# Digital Twin Brain Simulator: Harnessing Primate ECoG Data for Real-Time Consciousness Monitoring and Virtual Intervention

**DOI:** 10.1101/2024.05.17.594789

**Authors:** Yuta Takahashi, Hayato Idei, Misako Komatsu, Jun Tani, Hiroaki Tomita, Yuichi Yamashita

**Affiliations:** Department of Information Medicine, National Center of Neurology and Psychiatry, Tokyo, Japan; Department of Psychiatry, Graduate School of Medicine, Tohoku University, Sendai, Japan; Institution of Innovative Research, Tokyo Institute of Technology, Tokyo, Japan; Cognitive Neurorobotics Research Unit, Okinawa Institute of Science and Technology, Okinawa, Japan

**Keywords:** Digital twin, Treatment simulation, Real-time simulation, Artificial neural network, Generative artificial intelligence, Electrocorticogram, Personalized medicine

## Abstract

At the forefront of bridging computational brain modeling with personalized medicine, this study introduces a novel, real-time, electrocorticogram (ECoG) simulator based on the digital twin brain concept. Utilizing advanced data assimilation techniques, specifically a Variational Bayesian Recurrent Neural Network model with hierarchical latent units, the simulator dynamically predicts ECoG signals reflecting real-time brain latent states. By assimilating broad ECoG signals from Macaque monkeys across awake and anesthetized conditions, the model successfully updated its latent states in real-time, enhancing the precision of ECoG signal simulations. Behind the successful data assimilation, a self-organization of latent states in the model was observed, reflecting brain states and individuality. This self-organization facilitated simulation of virtual drug administration and uncovered functional networks underlying changes in brain function during anesthesia. These results show that the proposed model is not only capable of simulating brain signals in real-time with high accuracy, but is also useful for revealing underlying information processing dynamics.

## INTRODUCTION

In recent years, the concept of digital twins for human organs has emerged as groundbreaking technology that promises to revolutionize traditional medical frameworks [1-3]. A digital twin represents an organ’s functions through mathematical models, characterized by its ability to synchronize with the actual organ in real-time based on observational biological signals, thereby reflecting changes in the organ sequentially in the model [2, 3]. Such models, mirroring the current state of an individual’s organs, are anticipated to contribute to personalized treatment planning by enabling pre-emptive treatment simulations [1, 2]. Moreover, the capacity to precisely simulate effects of certain treatments holds promise for discovery of novel and more effective treatment approaches. Applications of digital twins for human organs have been progressively realized in fields such as cardiovascular disease treatment and pre-surgical simulations [1, 2]. In the fields of neurological and psychiatric disorders, significant heterogeneity in single disease populations underscores the heightened importance of personalized medicine, where benefits derived from digital twin technologies are expected to be substantial [4]. Until recently, the complexity of the brain was thought to pose significant challenges to construction of digital twins; however, advances in mathematical models for brain computations, the computational infrastructure for their implementation, and brain activity monitoring techniques have recently initiated embryonic efforts in development of digital twin brains.

Technological contributions to digital twin brain development have included creation of brain function simulators at various scales. At the microscopic scale, the focus has been on modeling components of neurons, including ion channels and receptors, as well as action potentials of individual neurons [5, 6] . These models offer the advantage of having a clear correspondence between neuronal properties, including receptors, and model variables, allowing for simulation of pharmacological effects. On the downside, modeling the whole brain based on microscopic neuronal models incurs immense computational costs [6]. There is also the difficulty of non-invasively acquiring microscopic data from patient brains, making it hard to incorporate real-time patient information. Furthermore, models at the microscopic level are still not methodologically mature enough to handle phenomena associated with higher-level cognitive processes. In contrast, a considerable volume of research has focused on modeling brain functions at a more macroscopic scale [7]. For instance, reinforcement learning models and Bayesian inference models represent the interaction between the agent and the environment with simple mathematical expressions, enabling modeling of cognitive processes as functions that output behavioral choices based on sensory input from the environment [7, 8]. However, the relationship between the limited number of variables that constitute these abstract models and the high-dimensional biological changes induced by therapeutic interventions, such as extensive changes in neuronal receptors and alterations in neural activity in various brain regions, remains unclear. Consequently, using these scale models for simulating therapeutic interventions still presents challenges.

In the context of bridging the gap between disparate scales of brain functionality, there has been a growing interest in recent years in simulations of brain functions at the mesoscopic level [9-12]. Unlike simulations that focus solely on individual neurons or local neural circuits, this mesoscopic level simulator entails modeling neural activities and their interactive dynamics on a regional basis in the brain. The mesoscopic level is where we can measure brain signals using non-invasive techniques like electroencephalography (EEG), magnetoencephalography (MEG), and functional magnetic resonance imaging (fMRI) to understand human brain function. It is also where we can apply neuromodulatory methods, such as transcranial magnetic stimulation and deep brain stimulation, showing how closely connected this level of modeling is to advanced medical technologies. Furthermore, simulations at the whole-brain level are feasible at manageable computational cost [13], making mesoscopic models promising candidates for digital twin brains. However, research on digital twin brains using mesoscopic models is still in nascent stages, and has yet to fully realize requirements such as real-time synchronization and interventional simulations. In this endeavor, our aim is to endow digital twin brains with requisite capabilities through utilization of generative artificial neural network models at the mesoscopic level. Our current model’s distinctive feature lies in its utilization of data-driven end-to-end training, an approach that autonomously extracts information from observed brain signals to accurately simulate brain functions responsible for generating these signals. To further enhance the model’s capacity for real-time, accurate simulations, “data assimilation” is employed. This advanced technique supports data-driven optimization even after training is complete, thereby enabling the model to dynamically adapt to unknown and changing situations. However, the inherent complexity and noise in brain signals present a substantial challenge. To address this, we employ stochastic generative models designed to capture intricate probabilistic structures underlying generation of these signals. This approach allows virtual therapeutic intervention to accurately simulate the process of generating changes in brain states and resulting brain signals.

In this study, we examine the utility of our proposed artificial neural network model as a digital twin brain, using electrocorticogram (ECoG) data. ECoG, one of intracranial EEGs, involves direct placement of electrodes on the brain’s surface, thereby offering higher precision in measuring brain activity than traditional EEG. By conducting data-driven training from ECoG signals, we aim to construct an ECoG generator (a digital twin brain prototype) and to evaluate its ability to meet the requirements of a digital twin, including real-time synchronization and intervention simulation, through the following procedure. Initially, we verify whether the simulator can generate ECoG signals with sufficient accuracy. Considering the clinical application of simulating novel patients after model development, the accuracy of generating ECoG for test individuals (not used for model training) is also verified. Furthermore, we evaluate the effectiveness of data assimilation, a technique to synchronize the digital twin with the brain in real time. This involves validating the model’s ability to progressively mirror the brain’s latent states, such as consciousness levels (awake and anesthetized states). Expanding on these findings, we conduct virtual intervention experiments on test individuals and assess the validity of the generated virtual ECoG. Additionally, by analyzing the functionality of this model, we endeavor to provide a unified explanation for the underlying generation system of anesthetic ECoG, a topic previously characterized by fragmented knowledge of frequencies and amplitudes and an incomplete understanding of its generation mechanisms. Specifically, by examining transmission of information to various brain regions in the model, we seek to define the functional network required for generation of anesthetic ECoG. Furthermore, through virtual intervention simulations on this functional network, we assess the contribution of each network to generation of anesthetic ECoG. Finally, we investigate whether the model can express individuality as a result of data-driven training and identify which variables represent this individuality.

## RESULTS

### System Overview

Development of the proposed digital twin brain simulator leveraged wide-area ECoG recordings of macaque monkeys, publicly accessible through the Neurotycho database [14, 15]. These data consist of ECoG signals recorded from four macaques, both under anesthesia (ketamine and medetomidine) and in awake, resting condition with eyes closed. Of the 128 channels covering an entire hemisphere of the cerebral cortex, ECoG signals from 20 channels spanning 10 brain regions were selected for use (Methods section for details). ECoG data of all four individuals were utilized, with three individuals used for training and the remaining individual used for testing (data assimilation), employing a four-fold cross-validation methodology.

The core component of the simulator is modeled using a Variational Bayesian Recurrent Neural Network (V-RNN), which is characterized by a temporal and spatial hierarchical structure (Figure 1A) [16-19]. This structure comprises three levels: the local region level (the first level) operating at short time scales at the lowest layer, the functional network level (the second level) functioning at intermediate time scales, and the global state level (the third level) that operates over long time scales at the highest layer. These level-specific dynamics are realized by latent units z, which represent the stochastic latent state of each level, and deterministic units d, which convey time-scale-specific deterministic dynamics.

**Figure 1.**
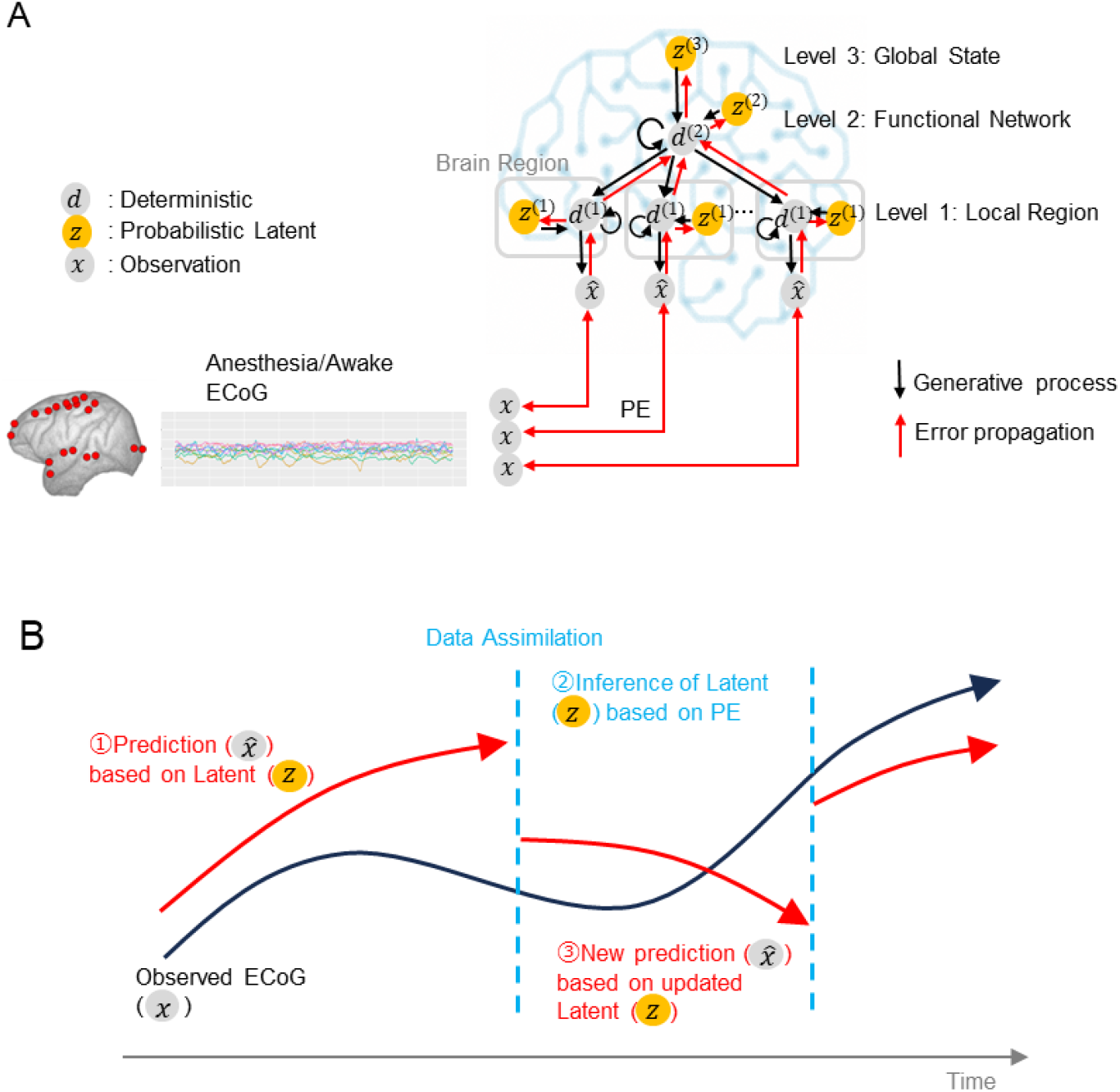
(A) Model Overview. This model depicts the process by which ECoG signals are generated from latent states *z* of the brain. The input *x* consists of wide-area ECoG measurements from 20 channels across 10 brain regions in macaques. The latent variable *z* possesses three hierarchical levels, each of which is stochastic. Latent states *z*, through synaptic connections of deterministic units *d*, lead to generation of ECoG signals 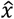. (B) Overview of the data assimilation procedure. For simplification, this figure does not show the time window. In an incrementally advancing window, predictions based on latent variables and updates of latent states based on prediction error (PE) are alternately repeated for a number of iterations.

The V-RNN operates according to a dynamic generative model acquired through data-driven training (Figure 1B), executing the following procedure: Initially, it estimates hierarchical and probabilistic latent states at time step *t* (*z*_*t*_), as the “prior.” Based on this prior of *z*_*t*_, the model predicts the ECoG signal 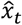. Subsequently, the actual observed ECoG signal *x*_*t*_ is inputted. The prediction error is calculated in a time window that encompasses the past spanning *H* time steps from *t* (*t* − *H* + 1: *t*). The latent state *z*_*t*−*H*+1:*t*_ is updated based on this prediction error and is referred to as the posterior. Predictions are then made again based upon this posterior. The operation of making predictions and updating the posterior is repeated a predetermined number of times. Finally, the posterior is used to generate the prior at the next time step (*z*_*t*+1_) through the deterministic state *d*_*t*_ . This sequential repetition of updating the latent state based on predictive and observational data, a process designated as data assimilation, enables real-time estimation of latent states and generation of virtual ECoG.

### Real-time latent state estimation via data assimilation

As a result of this training, the proposed model successfully utilized acquired internal dynamics to generate 20-channel ECoG sequences during both anesthetized and awake resting states. Supplementary Figure S1, which plots the loss throughout the training process, demonstrates a reduction in loss not only for training data, but also for validation data of test individuals. This indicates that the proposed model has acquired the capability to generalize, enabling it to generate ECoG sequences for test individuals.

Subsequently, the capacity of the proposed model to estimate underlying latent state changes of observed ECoG signals in real-time was evaluated using ECoG data that transitions from an awake state to an anesthetized state (or vice versa) in test individuals. Figure 2A presents an example of test results utilizing ECoG data during the transition from wakefulness to anesthesia. This figure plots observed and predicted ECoG signals (for 5 of 20 channels), their prediction errors, and the time series of latent states across the first to third levels (z^(1)^ to z^(3)^). A comparison between the observed and predicted ECoG waveforms reveals successful simulation of EcoGs by the model.

**Figure 2.**
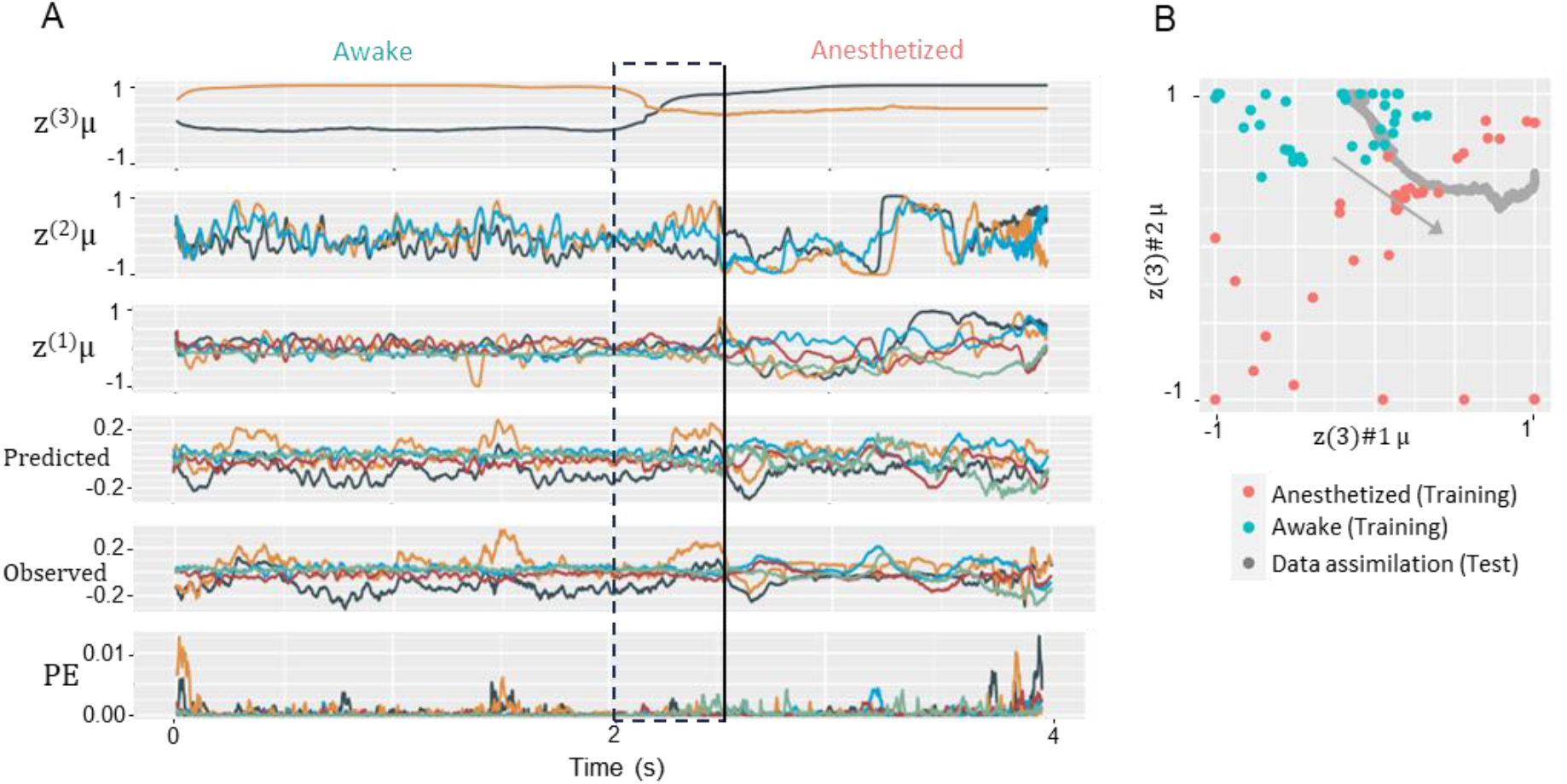
(A) Prediction and latent state sequences in data assimilation. This figure illustrates the sequence of predictions and the mean (μ) of posterior distributions of latent states z^(1)^, z^(2)^, z^(3)^, generated in real-time in response to observed ECoG data. The Observed ECoG combines 2500 time steps (2.5 seconds) of awake ECoG observations followed by 1500 time steps (1.5 seconds) of anesthetized ECoG observations from a test individual (an individual not included in the training dataset). This figure plots values after the data assimilation process is complete across all time steps of the observed ECoG sequence. The vertical solid line shows the moment of transition between awake and anesthetized conditions in observed ECoG data, with dashed lines showing the time window’s range when reaching this moment. In this range, updates of latent states are made in response to the transition in the observed ECoG. (B) Example of the distribution of the mean (μ) of z^(3)^ obtained during training and data assimilation phases. In this study, the latent unit z^(3)^ is prepared in two dimensions, and the x- and y-axes represent the z^(3)^μ values in the first and second dimensions, respectively. The gray arrow indicates the direction in which z^(3)^ changed over time during data assimilation.

Temporal changes in the latent state, reflecting data assimilation, are rapid at lower levels and slow at higher levels, depending on the time scale. Notably, at the highest level (z^(3)^), a switch in values can be seen in response to the transition from awake to anesthetized states (as enclosed in the box in Figure 2A), suggesting that the value of z^(3)^ indicates global changes in brain state between wakefulness and anesthesia. Dynamic alterations in these values throughout these processes are depicted in a Supplementary Video.

During data assimilation, changes in latent states z^(3)^ (two-dimensional) were plotted in gray on the scatter plot in Figure 2B. The red and cyan points represent the values of z^(3)^ for the anesthesia and awake conditions, respectively, as obtained through training. These observations revealed that clusters of latent states corresponding to anesthesia and awake conditions appeared to emerge. The trajectory of z^(3)^ across these clusters during data assimilation indicates that the model accurately estimated the transition from the brain states of anesthesia to wakefulness. To quantitatively validate this, data assimilation simulations were conducted using multiple ECoG sequences from test individuals, including transitions from wakefulness to anesthesia and vice versa. Using the silhouette width metric (Methods section for details), we evaluated whether the z^(3)^ states during data assimilation were appropriately positioned in clusters of wakefulness or anesthesia formed during training. These results showed that the proportion of correct cluster localization was, on average, 0.86 (SD 0.07) before the condition switch, and 0.73 (SD 0.17) after the switch. These findings suggest that the z^(3)^ states during generation of ECoG data for test individuals transitioning between wakefulness and anesthesia states can estimate these conditions.

### Virutual intervention to z^(3)^ (virtual drug administration)

The preceding analysis revealed that estimating latent states z^(3)^ from an test individual’s ECoG signals allows for prediction of anesthetic and awake conditions. Subsequently, we investigated whether virtual ECoG waveforms, generated by setting this estimated z^(3)^, accurately reflect characteristics of anesthetic and awake ECoGs. This approach, with z^(3)^ as a virtual intervention target, is intended to impact all hierarchical levels and brain regions downstream of z^(3)^, simulating administration of a drug that influences the whole brain. Evaluation of virtual ECoG leveraged a discriminator (3D convolutional neural network [20]) that was designed to differentiate between anesthetic and awake conditions (See Methods for details.). Results demonstrated that virtual ECoGs, generated by setting the estimated z^(3)^ for anesthetic and awake conditions in test individuals, were correctly classified into these respective conditions with high accuracies of 0.92 and 0.98. These results indicate that virtual ECoG signals generated by targeting z^(3)^ in virtual interventions accurately reflect the ECoG characteristics of both anesthetic and awake conditions.

### Functional Analysis of z^(2)^ (Functional network analysis)

We next explored representations that emerge in latent states of intermediate levels (z^(2)^) following training. The current V-RNN model incorporates multiple latent variables at the second levels (z^(2)^), each independently influencing deterministic units in the first level (d(1)) corresponding to various brain regions. Hence, each latent variable z^(2)^ is supposed to independently govern a distinct functional network. To analyze how these multiple functional networks change between anesthesia and wakefulness, we calculated transfer entropy as a measure to quantify the strength of control from z^(2)^s to d^(1)^s. Transfer entropy is a measure of the amount of information transfer between two sequences based on information theory.

In the current V-RNN, z^(2)^s incorporates three units, whereas d^(1)^s consists of modules that correspond to ten brain regions. Consequently, transfer entropy values from z^(2)^ to d^(1)^ in a single experiment are obtained as three sets of ten-dimensional vectors. Considering the implementation of cross-validation among four individuals, resulting in four separately trained networks, and given two conditions, anesthesia and wakefulness, it follows that 24 vectors (3 units × 4 networks × 2 conditions) were acquired. These vectors were subsequently sorted through hierarchical cluster analysis based on Euclidean distance (Figure 3A). Figure 3A reveals that transfer entropy values from z^(2)^ to d^(1)^ split into different clusters for anesthetized (top half of the heatmap) and awake (bottom half) conditions, with a tendency for higher transfer entropy under anesthesia. This indicates a stronger control mechanism from z^(2)^ to d^(1)^ under anesthesia than during wakefulness, reflecting the emergence of more explicit functional networks in the anesthetized state. Moreover, these transfer entropy values from z^(2)^ to d^(1)^ under anesthesia were composed of three clusters, as depicted in the dendrogram colored green, yellow, and red in Figure 3A. Each cluster has been designated as Functional Network (FN) 1, FN2, and FN3, respectively. These networks plausibly represent the necessary networks to generate ECoG during anesthesia. Figure 3B visualizes the average transfer entropy from z^(2)^ to d^(1)^ in FN1, FN2, and FN3 during anesthesia on the brain diagram, with increased transfer entropy observed in the frontal and anterior temporal areas for FN1, the frontal area and S1 for FN2, and the V1 for FN3. Figure 3C presents the transfer entropy during wakefulness for each FN, showing a general decrease in transfer entropy for FN1 and FN2 compared to the anesthetized condition, while V1 in FN3 remains high. These results demonstrate that the dynamics controlling each brain region vary between anesthesia and wakefulness, and these variations are distinct for each FN.

**Figure 3.**
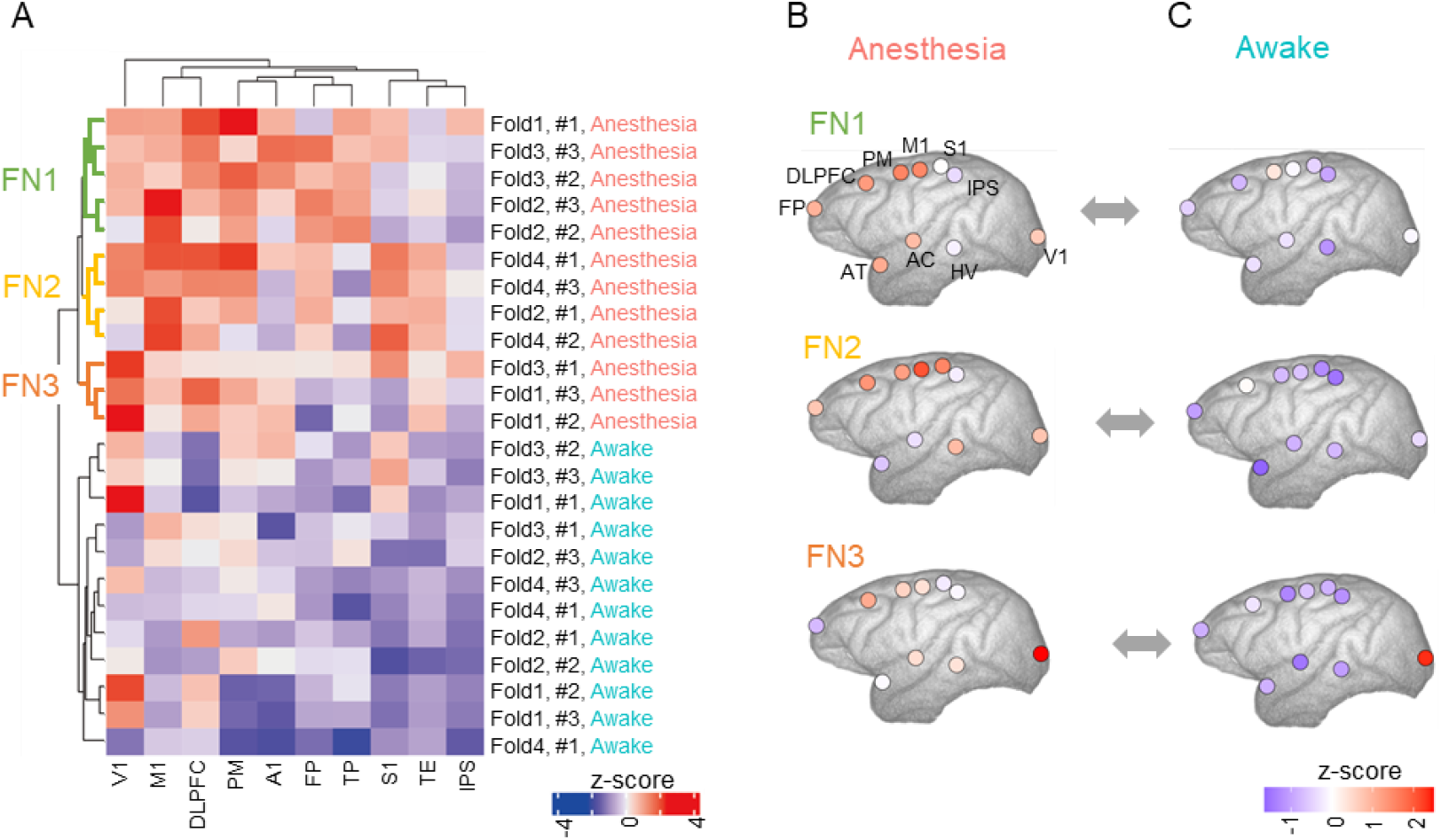
Transfer entropy analysis. (A) This heatmap represents transfer entropy values from z^(2)^ latent variables to d^(1)^ in various brain regions. Rows in the heatmap correspond to z^(2)^ latent variables used in calculation of transfer entropy. These rows represent 24 unique z^(2)^ states, derived from four variations in the training dataset during cross-validation (denoted as Fold 1-4), three z^(2)^ latent variables per one model (denoted as #1-3), and two conditions: anesthetized and awake. Columns indicate brain regions associated with d^(1)^ activities utilized in transfer entropy calculations. Hierarchical clustering was performed on rows and columns of this heatmap matrix. In this clustering, individual elements build a hierarchy by progressively merging clusters based on Euclidean distances. The heatmap color scale with transfer entropy values (z-score) from each z^(2)^ to the corresponding brain region d^(1)^, indicating the extent of control (or information flow). (B)(C) Mapping of transfer entropy values from z^(2)^ to d^(1)^ in different brain regions for FN1, FN2, and FN3. Abbreviations: FP, frontal pole; DLPFC, dorsolateral prefrontal cortex; PM, premortor cortex; M1, primary motor cortex; S1, primary somatosensory cortex; IPS, intraparietal sulcus; AT, anterior temporal cortex; AC, auditory cortex; HV, higher visual cortex; V1, primary visual cortex.

Subsequently, the impact of these FNs on altering ECoG patterns from wakefulness to anesthesia was investigated through virtual interventions targeting z^(2)^. For this virtual intervention, z^(3)^ states were set to those of an awake condition, under which ECoG signals characteristic of wakefulness would be generated in the absence of intervention. Thereafter, a virtual intervention was conducted on z^(2)^ states, altering each unit individually to a state representative of anesthesia. By applying the ECoG signals generated through this intervention to the discriminator described in the previous section, we can assess the extent to which generated ECoG signals transitioned toward those characteristic of anesthesia. This approach enables evaluation of causal effects of individual FNs on generation of anesthetic ECoG signals. These results intriguingly revealed significant variations in the proportion of anesthetized ECoG signals generated depending on the intervened FNs, with proportions of 0.12 (±0.06) for FN1, 0.74 (±0.25) for FN2, and 0.05 (±0.08) for FN3. These findings highlight the substantial differences in the causal effects of each FN to alter ECoG signals toward an anesthetic condition.

### Functional Analysis of z^(1)^

Similar to the scheme outlined in the previous section, we next investigated the impact of latent states at a local region level (z^(1)^) on altering generation of ECoG signals from awake to anesthetized conditions through a virtual intervention experiment. By setting z^(3)^s for the awake condition and z^(1)^ for the anesthetized condition in a targeted brain region, we generated ECoG signals and used the discriminator for condition differentiation. Table 1 presents the proportion of generated ECoG signals with those characteristic of anesthesia following intervention in specific brain regions. The average proportion of anesthetic ECoGs ranged from 19% to 29%, with a variance ≤10% and significant individual differences (SD). While the effect size was relatively small, there was a tendency for the proportion of anesthetic ECoGs to vary depending on the intervention brain region. Specifically, a two-way ANOVA model, with the proportion of anesthetic ECoGs as the dependent variable, and brain region and individual as independent variables, indicated a trend toward a significant main effect of the brain region on the proportion of anesthetic ECoGs, with a P-value of 0.064.

**Table 1.** Proportion of anesthetic ECoGs generated by intervention brain region

| Region | FP | DLPFC | PM | M1 | AT | AC | HV | S1 | IPS | V1 |
| --- | --- | --- | --- | --- | --- | --- | --- | --- | --- | --- |
| Mean | 0.21 | 0.23 | 0.24 | 0.19 | 0.22 | 0.26 | 0.27 | 0.28 | 0.29 | 0.21 |
| SD | 0.32 | 0.34 | 0.31 | 0.29 | 0.34 | 0.37 | 0.39 | 0.34 | 0.36 | 0.33 |

Abbreviations: SD, standard deviation; FP, frontal pole; DLPFC, dorsolateral prefrontal cortex; PM, premotor cortex; M1, primary motor cortex; S1, primary somatosensory cortex; IPS, intraparietal sulcus; AT, anterior temporal cortex; AC, auditory cortex; HV, higher visual cortex; V1, primary visual cortex.

### Individuality analysis

Finally, we assessed whether the proposed model could reflect individual characteristics, focusing on z^(3)^ representation. Figure 4A presents an example of the estimated probability density distribution of z^(3)^ states for awake and anesthetized conditions of the three individuals used in training. Figures 4BCD identify which individual corresponds to each point among these training results, while Figure 4E estimates the z^(3)^ states of an test individual. The distribution of z^(3)^ under anesthesia appears to vary among individuals, whereas the distribution under awake conditions appears to overlap. Quantitative evaluation based on cross-validation confirms individual-specific clustering under anesthesia (average silhouette width 0.228 > 0, P = 9.16×10^-11^), whereas no statistically significant cluster structure was observed under wakefulness (−0.09, P > 0.05). Thus, in terms of z^(3)^ representation, individual differences are self-organized under anesthesia, while individual representations are not distinctly observed under wakefulness.

**Figure 4.**
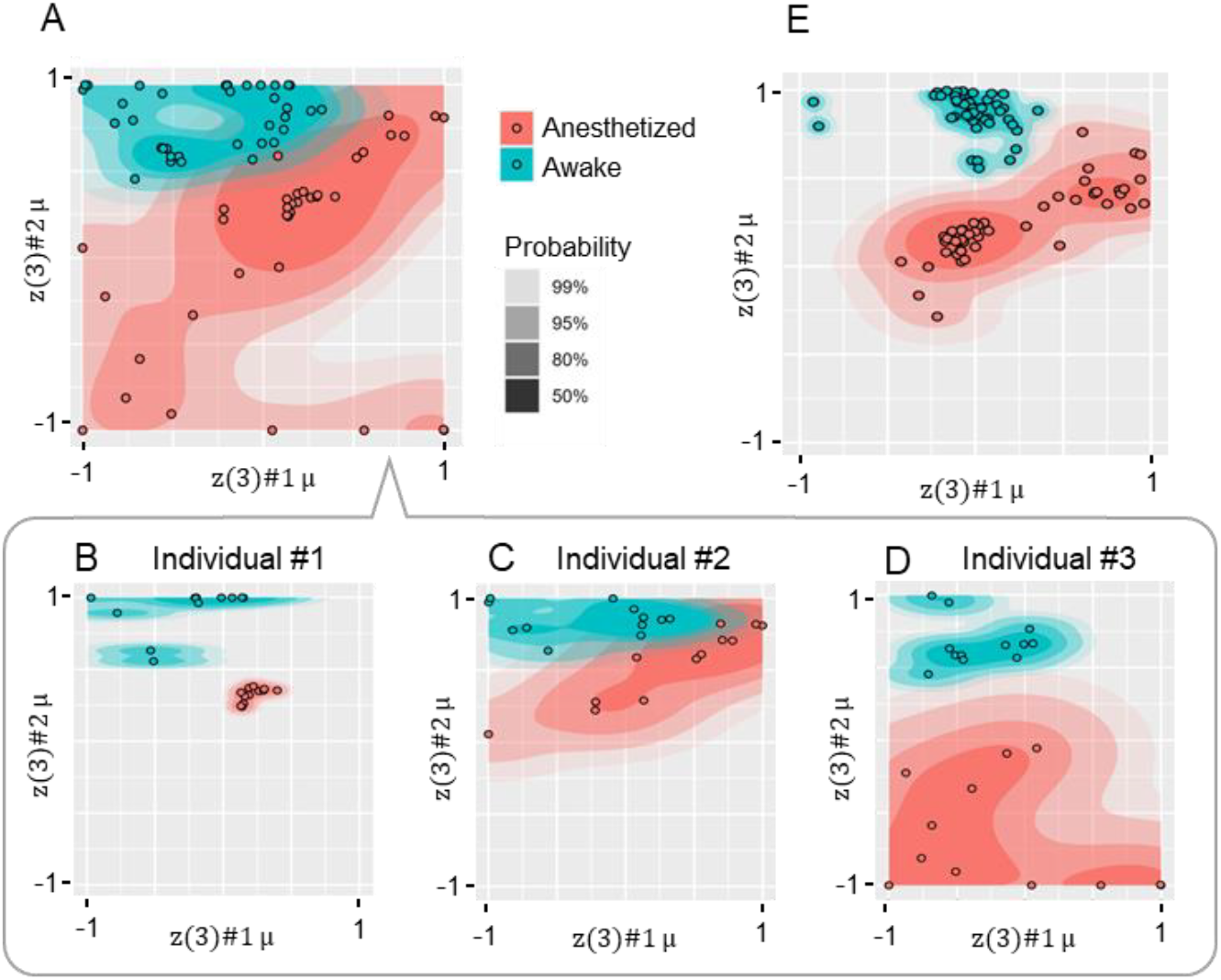
(A) Distribution of z^(3)^μ post-training, as shown in Figure 2B, overlaid with an estimated probability density plot. Each layer of varying shades represents boundaries of corresponding probability levels. Estimation of bivariate probability density utilized the kernel density estimator from the “ggdensity” package in R. (B) (C) (D) Distributions of z^(3)^μ depicted in Figure 3A are shown separately for each of three individuals in the training data. (E) Shows the distribution of z^(3)^μ estimations from data assimilation applied to ECoG sequences from a test individual.

Subsequently, we examined whether waveforms of virtual ECoGs generated based on the estimated z^(3)^ reflects individual characteristics. Evaluation of virtual ECoG leveraged a discriminator (3D convolutional neural network [20]) that was designed to distinguish among the four individuals used in the experiment (details in Methods). These results indicate that the proportion of correctly identified ECoG signals generated for test individuals was 0.35 under anesthesia and 0.30 when awake, both slightly exceeding chance levels (0.25 of selecting the correct individual out of four).

## DISCUSSION

### Summary of Results

In this study, we developed a digital twin brain prototype capable of generating ECoG signals from test individuals with high precision in real-time. This was achieved by employing a V-RNN with latent variables possessing temporal and spatial hierarchical structures, which updates these latent variables sequentially through data assimilation from observed ECoGs. Examination of the latent variable representation that facilitated such precise simulations revealed its self-organized structure at different hierarchical levels. Specifically, at higher levels of latent states (z^(3)^), clusters reflective of global brain states such as anesthesia and wakefulness, as well as individual uniqueness, were observed. Clusters of these z^(3)^ latent states enabled appropriate updates of latent states during data assimilation of ECoG signals from test individuals, leading to successful virtual drug administration simulations. Transfer entropy analysis between intermediate (z^(2)^) and lower (d^(1)^) hierarchical levels indicated emergence of activity patterns that control local brain regions corresponding to functional networks. Virtual interventions on these functional networks demonstrated that the effect of altering generated ECoGs from wakefulness to anesthesia varied significantly across functional networks.

### Comparison with conventional simulators

There have been efforts to achieve real-time simulation of ECoG and EEG signals. For example, in pursuit of EEG signal simulation, dynamics of average postsynaptic potentials (PSPs) in excitatory and inhibitory interneuron populations under electrodes have been modeled using differential equations [10, 21, 22]. These equations incorporate synaptic gain and time constants as time-varying parameters, with these parameters being inferred in real-time from observed EEG signals. Ultimately, the sum of the PSPs of excitatory and inhibitory interneurons is output as the prediction of the EEG signal. While this modeling approach is notably advantageous for its low computational cost, it is primarily designed for simulating a single-channel EEG signals [21], posing limitations on capturing intricate interactions among various brain regions. Concomitant with such real-time-focused model development, there has been an increase in attempts to create models that generate interactions among brain regions, reflecting individuality based on brain imaging [11, 12, 23, 24]. These efforts often employ graph network models, representing interactions among brain regions with differential equations that primarily involve linear transformations of state variables of brain regions. Parameters for these linear transformations, such as weights or distances of connections between brain regions, are determined based on diffusion brain MRI and related methods, which measures brain structures, allowing model individualization [23]. These models are frequently used to simulate the BOLD signal in fMRI. A notable feature of these models is that the model structure can be determined based on the actual brain structure of the individual, allowing discussion of the relationship between brain structure and brain function [12, 23]. On the other hand, technical challenges remain in realizing real-time data assimilation. A shared attribute among these models that are currently extensively utilized is a priori explicit formulation of dynamics specific to each brain signal type using differential equations [21, 23]. In contrast, the primary characteristic of our model using artificial neural networks is their data-driven training, offering substantial flexibility in handling input signals and facilitating modeling of diverse modalities, including simultaneous modeling of multiple signal types. In scenarios involving processing of brain signals, such as EEG, MEG, and fMRI, coupled with sensory input signals (visual and auditory), the proposed model’s general applicability could be valuable. On the other hand, reduction of computational cost in the real-time PSP simulator and incorporation of individual brain structures into the architecture of the graph network model are notable features of these previous models, suggesting that combining insights from these models can lead to refinement of the proposed model.

### Functional analysis using Digital Twin Brain

The proposed model not only achieves real-time, high-accuracy simulation of brain signals, but also has the potential to yield new insights into mechanisms of brain function changes during anesthesia, through functional analysis. Previous studies on ketamine, a primary component of the anesthesia used, have reported increased connectivity from subcortical areas, such as the thalamus, to the cortex [25], and decreased connectivity between the frontal and parietal regions [26, 27], particularly between M1 and S1 [28]. On the other hand, in the current study, we observed increased connectivity (transfer entropy) from z^(2)^ to local brain regions (d^(1)^) under anesthesia (Figure 3A) and found that FN2, which strongly controls brain regions M1 and S1, significantly impacts generation of anesthesia ECoGs. These findings suggest an association between z^(2)^ and subcortical functions. Moreover, decreased connectivity between the frontal and parietal regions during ketamine administration may be underpinned by enhanced control from hierarchical levels higher than these brain regions, exerted by the functional network, such as FN2. By this means, methodology employed in the current study could provide a new perspective, supplementing and explaining background processes behind the fragmented findings of prior research. Meanwhile, some findings of the current study seem to differ from previous research. For instance, while prior ketamine studies have identified reductions in connectivity from the frontal cortex not only to S1 but also to posterior parietal regions [27], the current study did not find much change in connectivity (transfer entropy) from z^(2)^ to the posterior parietal region (IPS) between anesthetized and awake conditions (Figure 3A). These seemingly divergent results may be attributed to differences in research methodologies. Whereas previous research focused on direct inter-regional information transmission, the current study assumed hierarchical latent states of top-down control over brain regions behind ECoG generation. Furthermore, unlike most prior studies, the current study implemented virtual interventions in functional networks to investigate their causal effects. We hope that approaches of previous studies and the current study complement each other, thereby contributing to a more comprehensive understanding of complex brain functions.

### Limitation

In this study, we conducted a virtual anesthetic ECoG generation experiment on test individuals to examine whether individual characteristics were replicated. As a result, it was confirmed that individual-specific clusters emerged in higher-order latent states z^(3)^; however, applying discriminators to generated ECoG signals yielded an identification rate of 0.34 for these individuals, which, although above chance level, remained relatively modest. Several factors may account for this outcome. First, generating ECoG signals for test individuals using only training data could be inherently difficult, as training data lack enough detail to accurately capture an test individuals’ unique characteristics. Second, optimization of network structural parameters (such as the number of neurons, hierarchical levels, and connectivity) might not have been adequate, suggesting that improvements in these parameters could enhance the capacity to replicate individuality. In addition to improve network structures, future studies are expected to explore how individuality is represented when increasing the number of training individuals or when dealing with human brain signals, instead of those of macaques. Furthermore, future research is expected to explore what the individual differences manifested in latent state z^(3)^ represent. In this study, clusters emerged only under the anesthesia condition and not under the awake condition. It cannot be ruled out that these individual differences under anesthesia represent a simple intervention intensity, such as the depth of anesthesia. Integrating biological signals other than ECoG or information about the intervention intensity into the model inputs may facilitate a deeper discussion on the relationship between intervention intensity and intrinsic individual differences.

### Brain computational theory and future perspectives

The proposed model can be discussed in relation to modeling of cognitive processes from the perspective of brain computation theory. The model executes free energy minimization through parameter optimization, which is a core aspect of the free energy principle derived from Bayesian brain theory [29, 30]. This theory hypothesizes that the brain’s information processing is analogous to data assimilation. Indeed, similar RNN models have been utilized in computational modeling and neurodevelopmental robotics research to understand cortical information processing, cognitive functions, and psychiatric symptoms [31-35]. The present research may be interpreted as demonstrating that the free energy principle-based approach is beneficial for simulating not only cognitive processes, but also brain signals. Future work is anticipated to expand this model to simultaneously simulate cognitive processes, including sensory inputs and motor outputs, in addition to brain signals, including ECoG, EEG and fMRI. This approach indicates the potential to uncover latent state changes underlying cognitive features in neuropsychiatric disorders and to facilitate virtual intervention experiments. Consequently, the model proposed here, with its ability to encapsulate multi-level phenomena, holds promise to illuminate brain information processing, potentially serving as a prototype for a digital twin brain.

## Methods

### Data and preprocessing

In this study, we utilized ECoG data from macaques, sourced from the publicly available Neurotycho database[14, 15, 36, 37]. To reduce computational costs, we selected a total of 20 channels, with two channels from each of the ten representative brain regions, from the entire set of 128 ECoG channels distributed across the hemisphere. These representative regions include the frontal pole (FP), primary visual cortex (V1), higher visual cortex (HV), auditory cortex (AC), primary somatosensory cortex (S1), primary motor cortex (M1), intraparietal sulcus (IPS), premotor cortex (PM), dorsolateral prefrontal cortex (DLPFC), and anterior temporal cortex (AT). ECoG signals were recorded at 1000 Hz. We conducted experiments using a cross-validation approach (training on three individuals, testing on one) with four individuals (George, Chibi, Kin2, Su), employing data collected during both under anesthesia (ketamine and medetomidine) and in an awake, resting condition with eyes closed.

Preprocessing of ECoG data encompassed re-referencing, outlier exclusion, and linear normalization. During the re-referencing process, the Common Median Reference method was applied. Outlier exclusion adhered to protocols from the preceding study using the same dataset [15], where data were divided into 2000 time-step (2-second) bins, and bins with values exceeding eight standard deviations, as well as their immediate neighbors, were excluded.

Consequently, 1.4% of data under anesthesia and 8.6% in awake macaques were excluded. For linear normalization, a scale transforming the range of values across all subjects, examination days, and experimental conditions to [-1, 1] was implemented.

Following these preprocessing steps, data were segmented again into 2000 time-step (2-second) intervals from which twelve anesthetized and twelve awake sequences were randomly selected from each of the three training individuals, totaling 72 sequences (144000 time steps,144 seconds) used for training. Then, to validate our model’s efficacy on test individuals’ ECoG data, we devised three sets of test data for different experiments. For the real-time latent state estimation experiment, we crafted 25 sequences by concatenating ECoG signals: the first 2500 time steps were drawn from randomly selected awake sequences, whereas the latter 1500 time steps were derived from randomly chosen anesthetized sequences. Another 25 sequences were similarly prepared, but with the order of states reversed, totaling 50 sequences. This disparity in time steps between the first and second halves accounts for consideration of a time window width *H* of 500 time steps. For the virtual intervention experiment to z^(3)^ and individuality analysis, we randomly selected 100 sequences of ECoG signals truncated at 2000 time steps, 50 under anesthesia and 50 while awake. Similarly, for virtual intervention experiments on z^(2)^ or z^(1)^, we randomly selected 50 sequences of ECoG signals truncated at 2000 time steps, 25 under anesthesia and 25 while awake.

### System Overview

This simulator’s main component employed a V-RNN, characterized by its spatial and temporal hierarchy. This hierarchy consists of the following three levels: a local region level (the first level), corresponding to modules for each of the 10 brain regions, operating on rapid changes at short time scales; the functional network level (the second level), consisting of a single module that controls local region modules and operates on slower changes over longer time scales; and the global state level (the third level), also a single module, which maintains constant latent states during generation of sequence predictions to map to sequential patterns of the target ECoG at an abstract level. These hierarchical structures are facilitated by each module comprising probabilistic latent units *z* and deterministic units *d* with time-scale-specific dynamics. In this model, generation of ECoG signals is characterized by top-down prediction, where information flows from latent units *z* to deterministic units *d*, descending to lower levels. Dynamics in these top-down predictions are acquired through a data-driven training phase. Following the training phase, a data assimilation phase utilizing ECoG data from test individuals facilitates real-time simulations. In the data assimilation process, initially, the latent states *z*_*t*_ at each hierarchical level for time step *t* are estimated as “priors”. Leveraging these priors, the model conducts top-down prediction of the ECoG signals 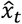. Subsequently, the actual observed ECoG signal *x*_*t*_ is inputted. Based on the prediction error (the difference between 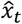 and *x*_*t*_), latent states *z*_*t*_ are updated, enabling a bottom-up estimation of the “posterior”. This posterior informs the next latent state *z*_*t*+1_ estimation, which becomes the new prior. This iterative process of top-down prediction and bottom-up posterior updates enables real-time estimation of latent states and the generation of virtual ECoG signals.

### Top-down prediction generation

In top-down prediction, the prior and posterior of latent states *z* are utilized to derive deterministic states *d*, enabling ECoG signal prediction 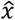. To illustrate this process, we initially discuss calculation of the prior for latent states *z*. Specifically, we consider the prior distribution 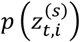 for the *i*th latent state 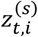 at time step *t* of the *s*th target sequence, at local region and functional network levels. This prior distribution is assumed to follow a Gaussian distribution, where the mean 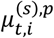 and standard deviation 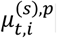 are derived from the previous deterministic states 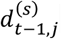.

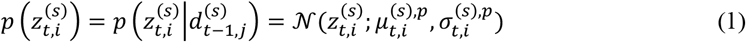

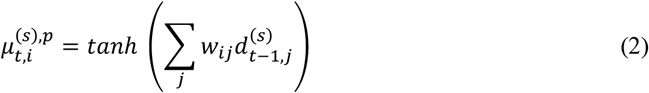

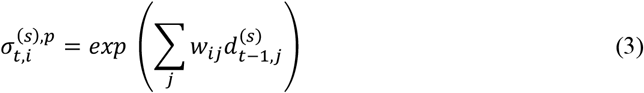

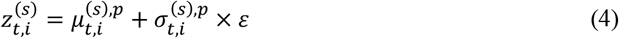

Here, *j* is an index representing deterministic units in the same module as 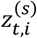. In Equation (4), *ε* is sampled from the *𝒩*(1,0) distribution to obtain the latent state 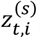. As described above, the latent state 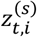 changes at each time step at local region and functional network levels, but at the global state level, the 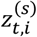 is kept constant during prediction generation. Therefore, at the global state level, 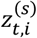 has a prior distribution of *𝒩*(1,0) only at the initial time step (*t*=1).

Next, the posterior distribution 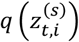 of the latent states is calculated as the conditional probability dependent on the prediction error signals 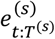.

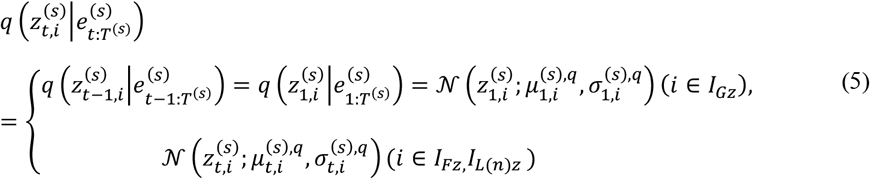

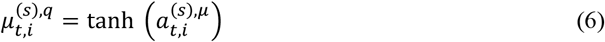

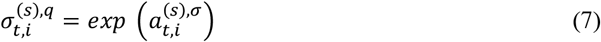

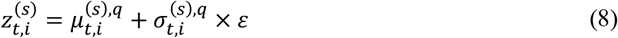

Here, *T*^(*s*)^ represents the length of the *s*th target ECoG sequence. 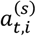 is defined as the unit’s adaptive internal state that represents the posterior distribution. 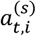 is dynamically updated at each timestep in every target ECoG sequence throughout the processes of learning and data assimilation. This updating process is influenced by the prediction error signal *e*_*t*:*T*_, which is propagated via a back-propagation-through-time (BPTT) algorithm. Thus, as inferred from equations (6) and (7), the posterior distribution of latent states may be conceptualized as a prediction-error-related unit state.

Based on calculated latent states 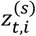 as described above, internal states 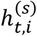 and outputs 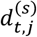 of deterministic units are computed as follows. This process involves deriving 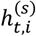 from the previous deterministic output 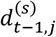 and latent states 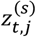 in the same module, alongside inputs from a higher-level module (Supplementary figure S2).

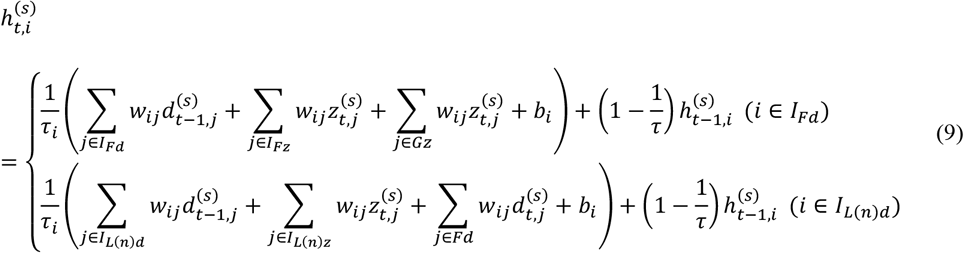

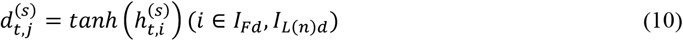

Here, *I*_*L*(*n*)*d*_, *I*_*Fd*_, *I*_*Gd*_ represent index sets for deterministic units at the local region level, functional network level, and global state level, respectively, while *I*_*L*(*n*)*z*_, *I*_*Fz*_, *I*_*Gz*_ represent index sets for latent units at their corresponding levels. The *L*(*n*) takes on *n* = 1,2, …, 10, corresponding to modules representing ten local brain regions. The *w*_*ij*_ refers to the weight of synaptic connections between units, and *τ*_*i*_ denotes the time constant. Deterministic units with smaller time constants *τ*_*i*_ tend to exhibit rapid changes in states, whereas units with larger time constants *τ*_*i*_ show a tendency for slower changes in states[19]. By setting *τ*_*i*_ = 2 at the local region level and *τ*_*i*_ = 4 at functional network level, the model represents changes on a shorter timescale at the lower level and on a longer timescale at the higher level, effectively constructing a temporal hierarchical structure.

Lastly, the prediction 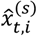 of ECoG signals for 20 channels corresponding to 10 brain regions is generated by the respective local brain region module.

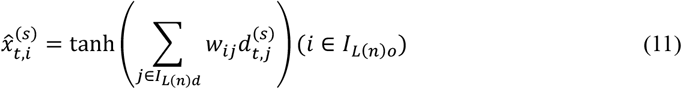

Here, the *I*_*L*(*n*)*o*_ denotes index sets of output units in the brain region *L*(*n*).

### Parameter updates through bottom-up optimization

Optimization processes in both training and data assimilation phases have been implemented by employing the negative Evidence Lower Bound (ELBO) as the loss function. This negative ELBO is equivalent to the variational free energy [38] in the context of brain computation theory, and is hereafter identified as *F*_*t*_. In the framework of V-RNN, *F*_*t*_ is derived as follows[18].

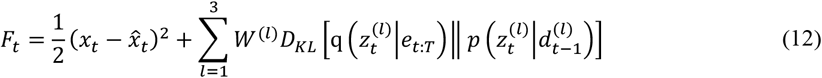

The first term of this equation represents the prediction error, while the second term denotes the influence of the prior 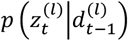 on the update of the posterior 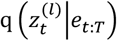 of the latent state. Consequently, in an optimization process aimed at minimizing *F*_*t*_, the update of the posterior progresses in a manner that minimizes the prediction error under the influence of the prior. The hyperparameter *W*^(*l*)^, referred to as the meta-prior[17], serves to balance the prediction error and the divergence between the prior and posterior, and can be individually set for each hierarchical level *l*. In this study, we set the meta-prior to a uniform value across all levels (*W*^(1)^ = *W*^(2)^ = *W*^(3)^ = 0.001). This optimization by minimizing *F*_*t*_ was carried out during both the training and data assimilation phases.

During the training phase, synaptic weights *w* and the posterior are updated to minimize the free energy 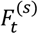 over all time steps and target sequences.

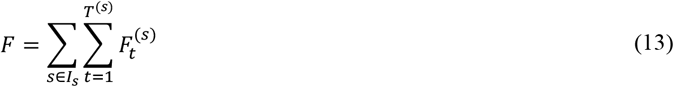

On the other hand, during the data assimilation phase, only the posterior is updated, while the weights *w* remain fixed. In this phase, the free energy is summed over the time window spanning the current time step *t* backward through *H* time steps.

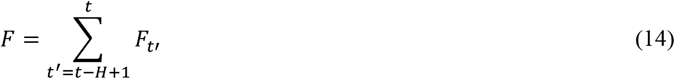

Utilizing the summed free-energy, the posterior of *z*_*t*−*H*+1:*t*_ in time windows of all modules is updated, with this window advancing over time. In both training and data assimilation phases, the Rectified Adam optimizer was employed for parameter updates [39]. Partial derivatives of the free energy with respect to each optimization parameter were computed using the BPTT algorithm.

### Data assimilation

Following the training phase, the data assimilation process is applied to ECoG signals of test individuals. Data assimilation involves a cyclical process of bottom-up posterior updates informed by top-down predictions and observations. This process occurs in a sliding time window that spans from the current time step *t* backward through *H* time steps, and incrementally moves forward as time progresses (Supplementary Figure S3). The process unfolds as follows: (1) The prior of *z*_*t*_ is estimated based on the posterior of the previous time step. (2) Based on this prior of *z*_*t*_ and the posterior at earlier time steps (*z*_*z*−*H*+1:*t*−1_), ECoG signals 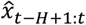 in the time window are predicted in a top-down manner. (3) Subsequently, new observed ECoG signals *x*_*t*_ are inputted. (4) At each time step in the time window, the free-energy *F* is calculated based on the prediction error (the discrepancy between *x* and 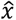). (5) To minimize the free-energy *F*, latent states *z*_*t*−*H*+1:*t*_ are updated as the posterior. (6) Based on this updated posterior, top-down predictions for 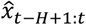 are performed again. This process of bottom-up posterior updates (4)(5) and top-down prediction (6) is repeated a predetermined number of times. Subsequently, the process returns to (1), where, from the posterior of *z*_*t*_, the prior for the next time step *z*_*t*+1_ is estimated. Such data assimilation enables real-time estimation of latent states and generation of virtual ECoG signals.

### Parameter setting

Based on preliminary investigations, the number of latent units *z* and deterministic units *d* have been configured to accurately predict ECoG signals. At the local region level, for each module in a respective brain region, one *z* and fifteen *d* units are allocated. At the functional network level, three *z* and fifteen *d* units; at the global state level, two *z* units. Note that at the global state level, the *z* states are set to remain constant during prediction generation and *d* units are not used. During the training phase, synaptic weights *w* and the posterior were updated 200,000 times. In the data assimilation phase, specifically during real-time latent state estimation experiments, the time window width *H* was set to 500 time steps, with the posterior updated 100 times at each timestep. For virtual intervention experiments, the time window width *H* was fixed to the sequence length (2000 time steps), and the posterior was updated 10,000 times. The reason for fixing time window to sequence length here is to obtain a representation of z^(3)^ that abstractly corresponds to characteristics of the full ECoG sequence length by not allowing z^(3)^ to change temporally. Hyperparameters of Rectified Adams during learning and data assimilation phases were set with α = 0.001 (learning rate), β1 = 0.9, and β2 = 0.999.

### Virtual intervention

Virtual intervention experiments were performed on latent states of z^(3)^, z^(2)^, z^(1)^. In these experiments, we generated virtual ECoGs by inputting latent states estimated as anesthetized or awake test individuals in target latent units. Then, we investigated whether resulting virtual ECoG waveforms exhibited characteristics of anesthetized or awake conditions. Latent states used as intervention inputs were posterior distributions obtained through data assimilation using ECoG signals of test individuals (See also the section of Parameter setting).

In virtual interventions targeting z^(3)^, the posterior of the constant z^(3)^ was input to latent units together with the posterior of z^(2)^ and z^(1)^ to generate virtual ECoGs. Effects of this virtual intervention were evaluated based on the proportion of generated ECoGs indicating anesthesia or awake characteristics. Assessment of these characteristics in ECoG signals utilized a discriminator, as described below.

As part of functional analysis, virtual intervention targeting z^(2)^ and z^(1)^ were conducted. This was undertaken to investigate the contribution of each latent unit in altering generated ECoGs from awake to anesthetized conditions. Initially, the posterior of an awake condition was input to z^(3)^ so that awake ECoGs were generated in the absence of intervention. Subsequently, the posterior of anesthesia condition was individually input to specific latent units of z^(2)^ or z^(1)^ (virtual intervention) and virtual ECoGs were generated. The contribution of each latent unit to anesthetized ECoG generation was evaluated based on the proportion of generated ECoG manifesting anesthetic characteristics, which was assessed by the discriminator, as described below. In these analyses, target latent units had their posterior distributions input, while non-target latent units underwent top-down prediction generation using prior distributions without explicit setting of distributions corresponding to awake or anesthetized conditions.

### Discriminator

In order to examine the characteristics of virtually generated ECoG signals, a discriminator was used. This discriminator made classifications as to whether the virtual ECoG signals pertained to states of anesthesia or wakefulness, or in which of the four individuals ECoG signals originated. The discriminator employed a model that converts ECoGs into spectrograms and utilizes a deep convolutional neural network (CNN) for classification, noted for its high accuracy in previous studies[40]. Training for this model utilized ECoG signals across all individuals in the study, during both anesthesia and wakefulness. Generation of spectrograms was carried out by applying the multi-taper time-frequency spectrum method to ECoG sequences that were segmented every 2000ms. During this process, the length of the moving window was set to 100ms, with a step size of 10ms, and calculations were performed for the 0-200Hz frequency band. Spectrogram images across 20 channels were compiled as a single-data unit and subjected to discriminant analysis using a three-dimensional CNN, the structure of which is detailed in Supplementary Figure S4. In the training phase of this CNN, cross-entropy was employed as the loss function, and the model was updated 100 times using an Adam optimizer with a learning rate of 0.0005. Models were constructed for both binary classification tasks, distinguishing between anesthesia and wakefulness, and for a four-class identification task involving individual differentiation. Test accuracies for these models were remarkably high, at 99.6% and 98.1%, respectively.

### Silhouette analysis

The cluster structures emerging in the distribution of latent states *z*(3) at a global state level were quantitatively assessed. For this assessment, silhouette width [41], a method commonly used for cluster evaluation, was used for the *z*(3)μ space as shown in Figure 3A, and the calculation process is shown below. Consider the case where data point *i* corresponding to one *z*(3)μ state (corresponding to one point in Figure 3A) is to be evaluated as to whether it belongs to cluster *C*_*i*_ . To begin with, a(*i*) is defined as follows.

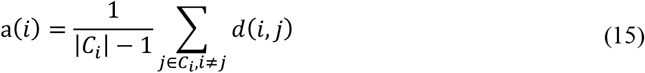

Here, |*C*_*i*_| is the number of points belonging to cluster *C*_*i*_, and *d*(*i, j*) is the distance between data points *i* and *j*. a(*i*) represents the average of the distances from data point *i* to the rest of the points in its cluster *C*_*i*_, indicating that a reduced a(*i*) value reflects greater cohesion in cluster *C*_*i*_ . Next, b(*i*) is defined as follows.

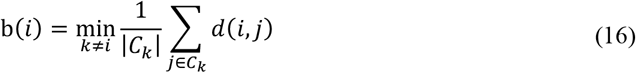

Here, b(*i*) represents the average distance between data point *i* and points in the adjacent, but distinct cluster, indicating that a higher b(*i*) value suggests a greater separation from adjacent clusters.

Based on a(*i*) and b(*i*), the silhouette width s(*i*) for data point *i* is calculated as follows.

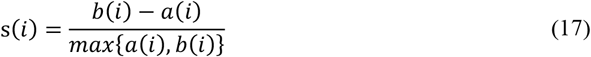

A higher value of s(i) indicates that data point *i* is more strongly clustered in *C*_*i*_ . In real-time latent state estimation experiments, we utilized s(*i*) to classify each time step’s *z*(3)*μ*, obtained through data assimilation, as belonging to either the anesthesia or awake clusters established during the learning process. Specifically, the s(*i*) was calculated for both the anesthesia and awake clusters, and data *i* was classified into the cluster with larger s(*i*).

From equation (17), it can be inferred that if s(*i*)> 0, data point *i* tends to be clustered into *C*_*i*_, rather than being randomly distributed. In this study, we utilized this principle to investigate whether individual clusters emerge in the *z*(3) space. Specifically, the individual to which each data point *i* belongs were designated as *C*_*i*_ and s(*i*) was calculated for all data points in the training ECoG sequences to statistically examine whether the average silhouette width exceeds zero (t-test).

## Data Availablity

ECoG data utilized in this study is available at: http://www.neurotycho.org/

## Code Availability

The code for the V-RNN, a principal component of the model discussed herein, is shared at: https://github.com/h-idei/pvrnn_sa

## Acknowledgements

This work was supported by JSPS KAKENHI (JP21K15723, JP24K20897, JP22KJ3167, JP20H00625, JP24H00076, JP24K00499, JP24H02175), JST CREST (JPMJCR21P4), JST Moonshot R&D (JPMJMS2031), AMED (JP21tm0424601), and Intramural Research Grant (4-6, 6-9) for Neurological and Psychiatric Disorders of NCNP.

## Author contribution statement

YT and YY conceived the study, HI contributed to model building, YT conducted experiments and analysis, YT prepared the original draft, and YT, HI, MK, JT, HT and YY conducted peer review and editing.

## Supplementary Figures

**Supplementary Figure S1:**
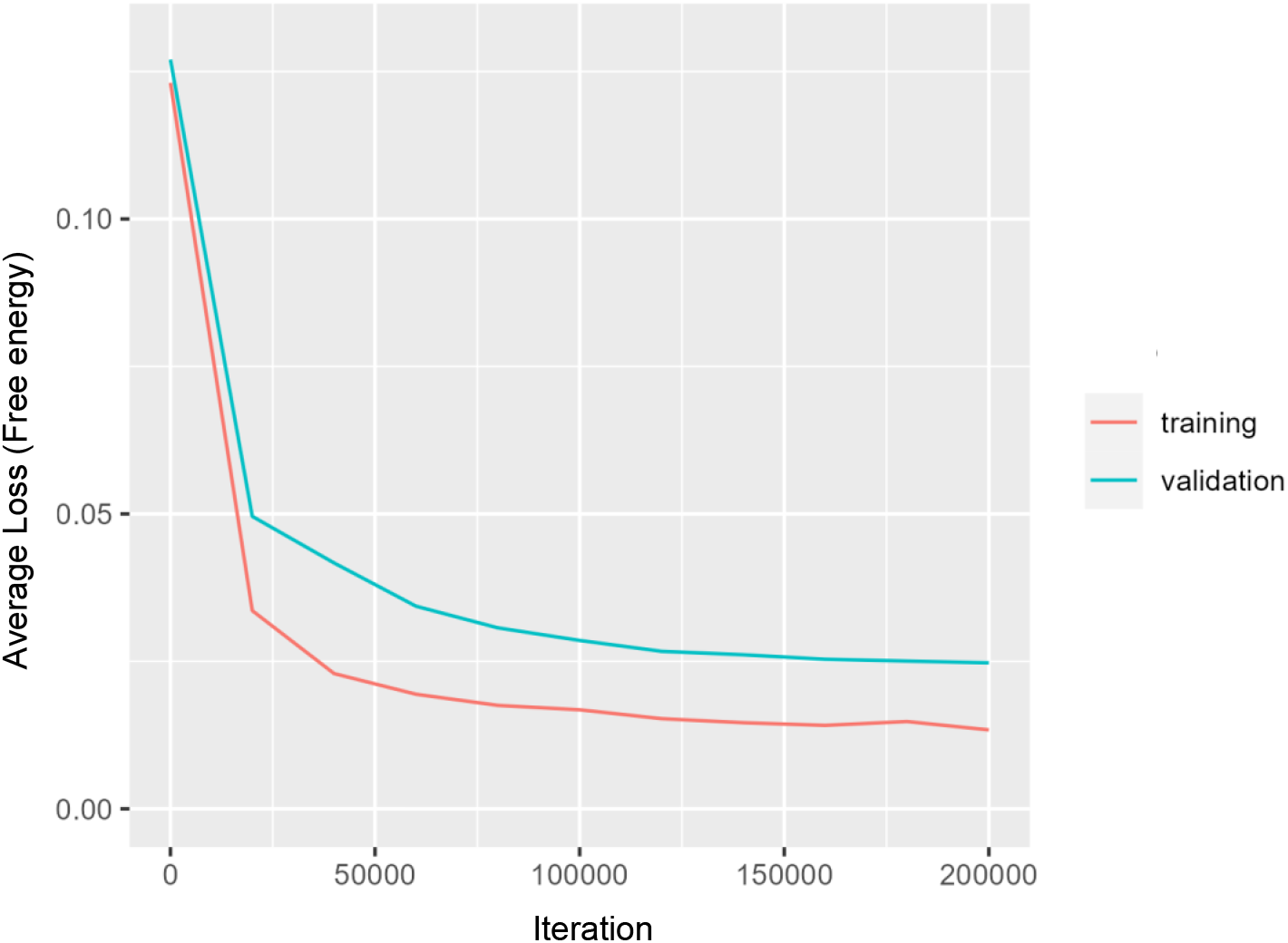
Learning curve.

**Supplementary Figure S2:**
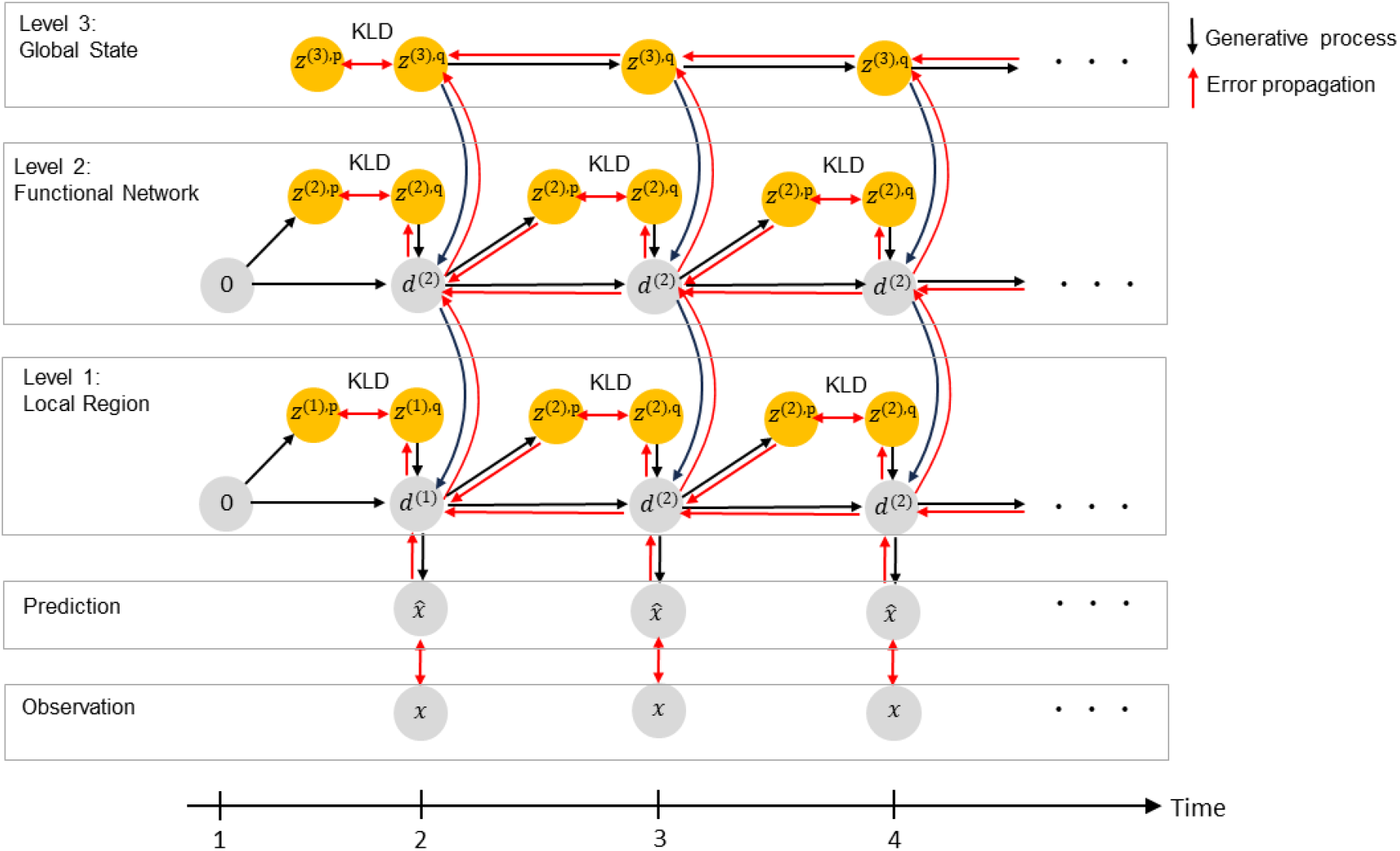
Temporal processing of the V-RNN. During the training phase, the network optimizes the posteriors of latent variables *z*_1:*T*_ across all modules and updates time-constant synaptic weights by minimizing the free energy F over the duration *T* of training ECoG sequences. Initial deterministic states of all modules are set to zero. For simplicity, only one out of ten modules is depicted for local region modules. Abbreviation: KLD, Kullback-Leibler divergence.

**Supplementary Figure S3:**
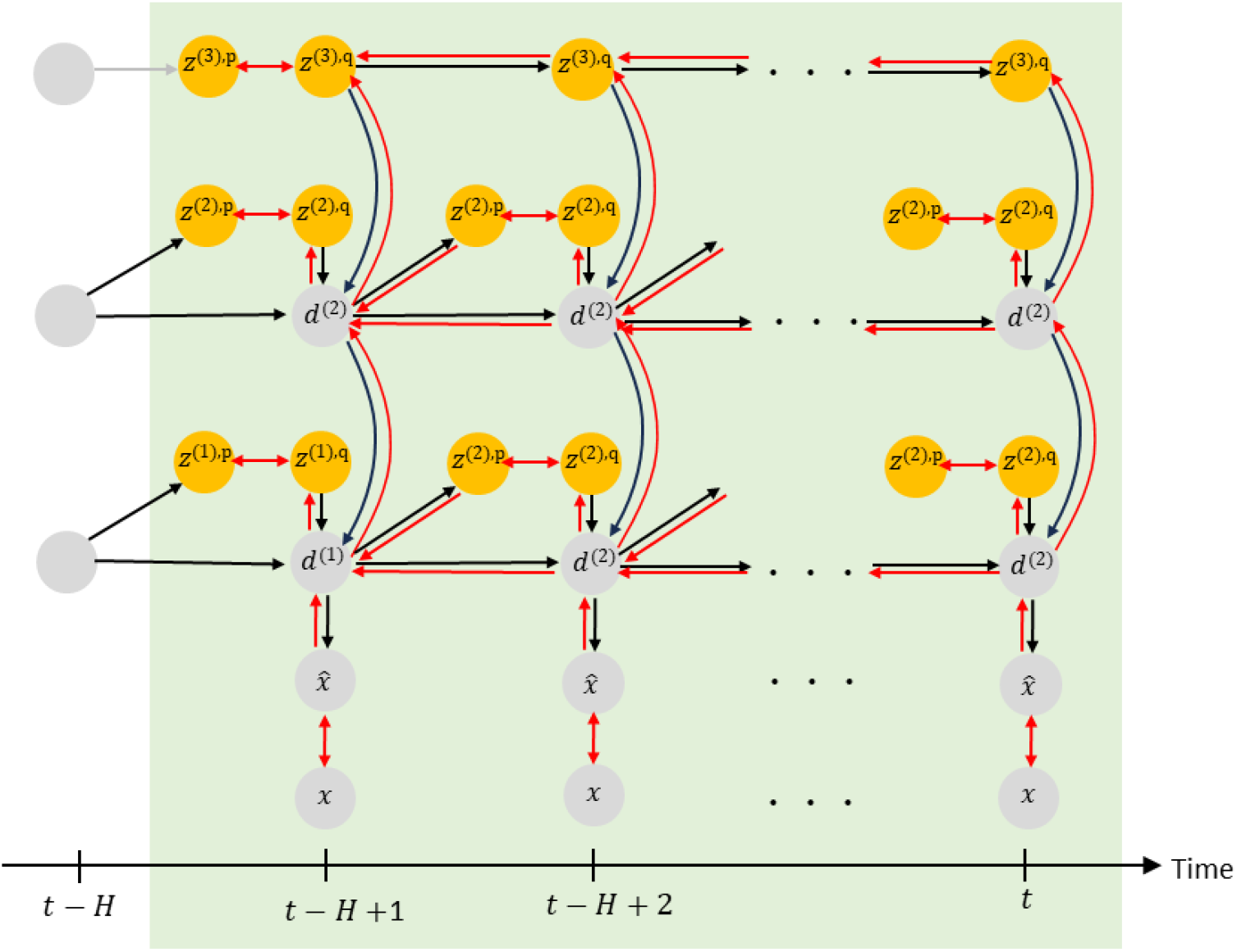
Data assimilation phase Data assimilation encompasses a cyclical process that integrates top-down predictions with bottom-up posterior updates driven by observations. This iterative procedure is conducted in a sliding time window that retraces *H* time steps from the current time step *t*, progressively moving forward as time advances. During this phase, synaptic weights remain fixed.

**Supplementary Figure S4:**
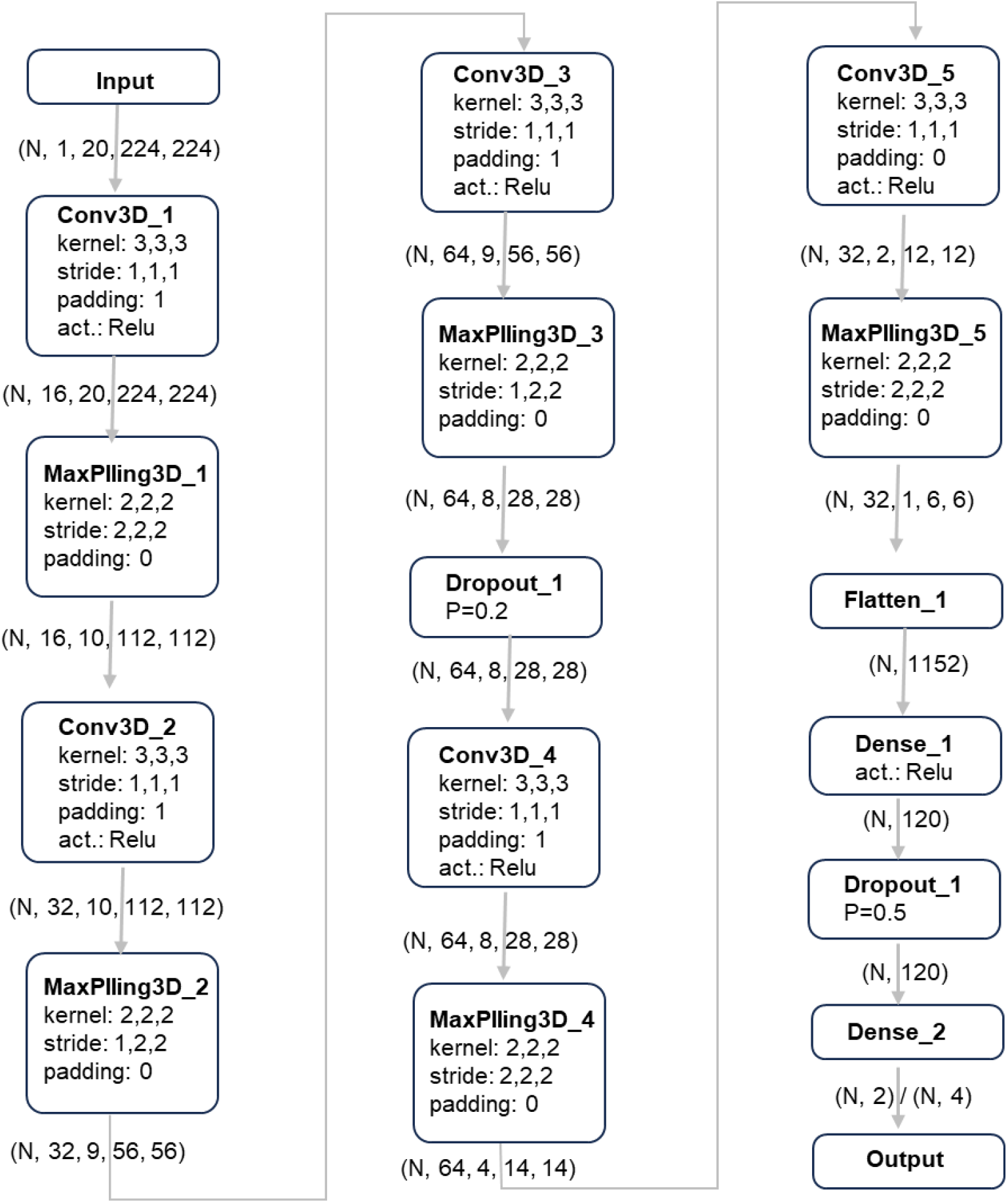
Network architecture of discriminator (3D convolutional neural network).

