## Supplementary figures and images for "Digital Twin Brain Simulator: Harnessing Primate ECoG Data for Real-Time Consciousness Monitoring and Virtual Intervention"

### Supplementary Video

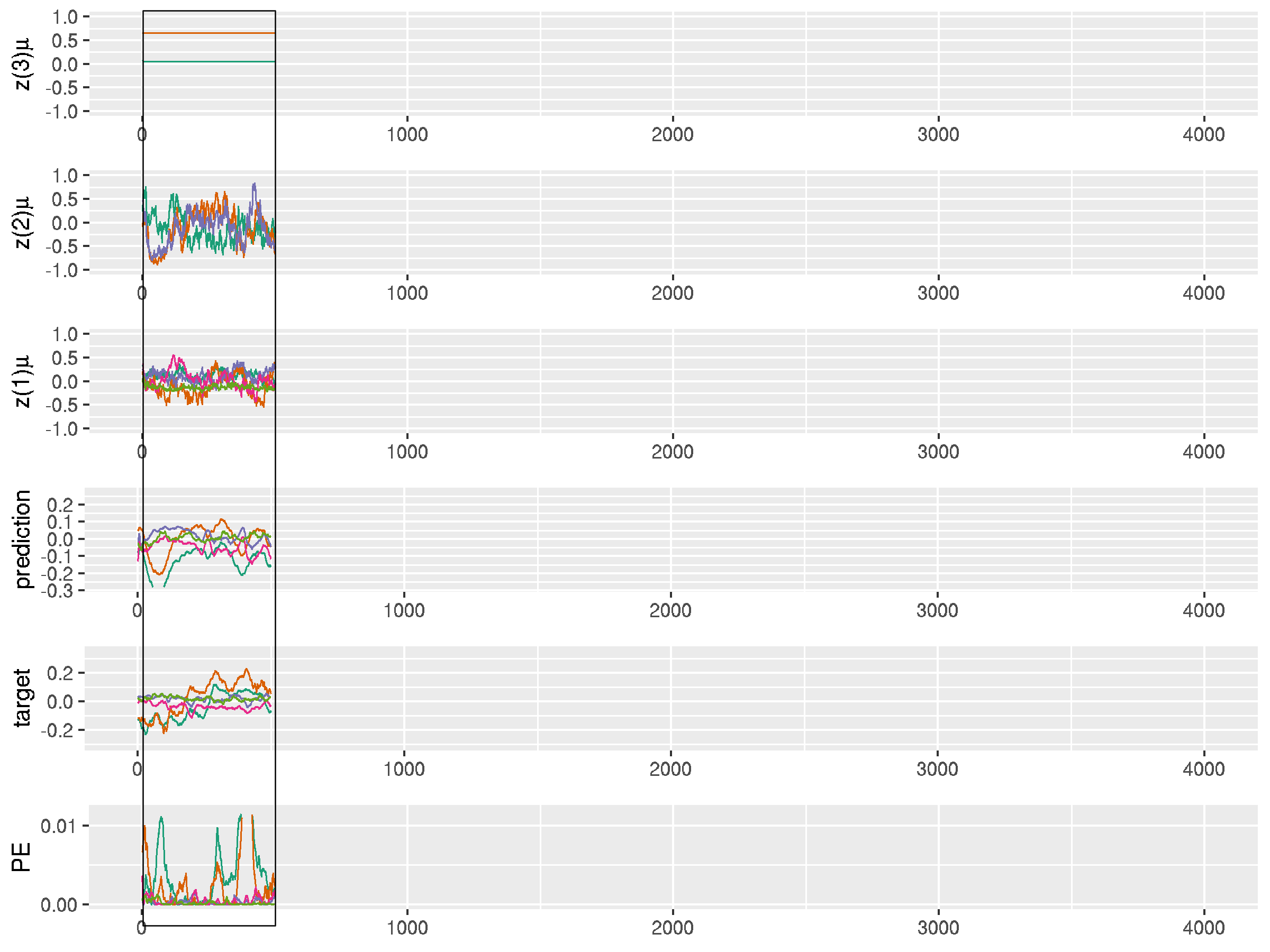
